# Exploratory Proteomics of Brain Extracellular Vesicle Enriched Preparations After LPS Challenge and miRNA Associated Vesicle Administration

**DOI:** 10.64898/2026.10.06.757102

**Authors:** Kent Brown, Yingjie Wang, Zhiying Shan, Lanrong Bi

## Abstract

We used brain extracellular vesicle (EV) proteomics to identify protein responses to acute neuroinflammation and an EV-based microRNA intervention. Rats received vehicle, intracerebroventricular lipopolysaccharide (LPS), or LPS followed by an intravenous EV formulation containing anti-inflammatory microRNAs (miRNAs). Brains were collected 24 h after LPS or vehicle administration, and three independently processed animal-derived EV-enriched preparations per group were analyzed by label-free proteomics. Across all nine samples, 953 protein groups had complete normalized abundance measurements. Compared with vehicle controls, LPS exposure was associated with 227 exploratory protein changes, comprising 95 increases and 132 decreases at nominal P < 0.05 and an absolute log_2_ fold change ≥ log_2_(1.5). Increased proteins were enriched for translation-related annotations, whereas decreased proteins were enriched for plasma-membrane annotations. These LPS-associated changes did not remain significant after Benjamini-Hochberg correction at a false discovery rate (FDR) threshold of q < 0.05. Following treatment, 119 of the 227 candidates were closer to control abundance on the log_2_ scale; 23 candidates also showed nominal treatment-versus-LPS differences. Lamb2 and Ptgfrn were the two proteins that differed significantly between treated and LPS-only animals after FDR correction, identifying them as priorities for further validation. Nevertheless, 174 proteins differed significantly between treated animals and vehicle controls, and analysis across the measured proteins did not demonstrate an overall shift toward control values. Thus, the intervention was associated with specific protein responses, but treated animals retained substantial differences from controls. This study identifies candidate markers for investigating treatment responses in brain EV-enriched preparations and demonstrates the value of examining individual proteins alongside broader proteomic changes. Independent validation is needed to determine their functional relevance and distinguish miRNA-specific effects from contributions of the administered EV carrier.

## 1. Introduction

Neuroinflammation changes the molecular environment shared by neural, glial and vascular cells. Extracellular vesicles (EVs) are one component of this environment, and their measured protein composition can vary with the condition of the source tissue. In cultured microglia, polarization produces distinct EV-associated molecular profiles, and EV preparations from LPS-activated cells can induce inflammatory transcriptional changes in recipient cells [1]. In mice, acute LPS exposure also alters the protein composition of brain-derived EV preparations, with changes that differ from those measured in bulk tissue [2]. These observations motivate analysis of EV-enriched fractions as potential indicators of inflammatory state.

An intervention introduces a second question: does the recovered protein profile approach the control state, retain the LPS-associated pattern, or acquire additional treatment-associated differences? These possibilities can coexist. Movement opposite to an LPS-associated change is a useful descriptive observation, but it does not by itself show restoration of normal abundance or function. A protein may overshoot the control value, and the overall profile may remain different from control despite selected reversals. Three time-matched groups are therefore useful for separating the LPS association, the treatment association, and the residual difference from control.

Two limitations require particular attention. First, tissue-derived EV preparations may contain nonvesicular particles and intracellular material released during tissue processing. In accordance with MISEV2023, operational descriptions and direct characterization are needed before attributing proteins to a particular EV subtype or cellular origin [3]. Second, a small discovery cohort provides uncertain estimates of protein-level variation. Nominal screening thresholds can prioritize candidates, but they do not replace false-discovery control or independent validation [4].

Here, we profiled brain EV-enriched preparations from vehicle, LPS, and LPS plus miRNA-associated EV groups collected at the same 24 h endpoint. We combined replicate-level abundance analysis with corrected pairwise testing, confidence intervals, explicit descriptions of movement toward control, and sensitivity analyses. We also tested the broad functional annotations supplied with the proteomic export against the quantified-protein background. The objective was to define a reproducible set of treatment-associated observations and validation candidates without inferring an unmeasured mechanism of EV loading or therapeutic action.

## 2 Materials and methods

### 2.1 Animals and experimental design

Adult male Sprague-Dawley rats (8-12 weeks; 200-350 g) were housed under standard conditions with food and water available ad libitum. Procedures were approved by the Institutional Animal Care and Use Committee. Three groups were studied: vehicle control (SD), LPS, and LPS plus miRNA-associated EVs (Mi). Each group comprised three animals, each contributing one independently processed brain preparation; samples were not pooled.

LPS from Escherichia coli O26:B6 (Thermo Fisher Scientific) was administered by a single unilateral intracerebroventricular injection of 5 µg in 2 µL into the lateral ventricle (AP -0.8 mm, ML +1.4 mm, DV -3.5 to -3.7 mm from the skull surface). Injection took approximately 2 min, followed by a 5 min needle dwell. Controls received 2 µL saline at the same coordinates. After recovery, the Mi group received a single tail-vein injection of the miRNA-associated EV preparation (20 µL per rat). All brains were collected 24 h after LPS or vehicle administration. No unloaded-EV or formulation-reagent control arm was included.

### 2.2 Preparation of the administered miRNA associated EV formulation

EVs isolated from untreated Sprague–Dawley brain were combined with three mature miRNAs [hsa-miR-146a-5p, hsa-miR-10a-5p, hsa-miR-21-5p, ThermoFisher Scientific] using Exo-Fect (System Biosciences). Each reaction contained 10 pmol of each miRNA, 15 µL Exo-Fect, 30 µL miRNA mixture, EV material equivalent to 300-500 µg protein, and sterile PBS to 225 µL. The mixture was inverted, incubated at 37 °C with shaking for 10 min, and placed on ice. ExoQuick-TC (45 µL) was added, followed by mixing and incubation on ice for 30 min. Material was pelleted at 13,000-14,000 rpm for 3 min and resuspended in 120 µL PBS; each animal received 20 µL.

The administered material is described as a miRNA-associated EV formulation. The preparation procedure alone does not establish whether miRNA is intraluminal, surface-associated, reagent-associated, or distributed among these states. Treatment-associated differences are attributed to the complete formulation rather than to the miRNAs alone.

### 2.3 Brain EV enrichment and characterization

The whole brain, excluding the cerebellum, was used for EV isolation following a protocol described previously [22]. Briefly, brain tissue was cut into small pieces and incubated in Gibco Hibernate E medium at a ratio of ∼0.2 g tissue per 2 mL medium in each well of a 6-well plate. Collagenase D and DNase I were added to final concentrations of 2 mg/mL and 40 U/mL, respectively, and the tissue was incubated at 37 °C for 30 min with gentle shaking at 70 rpm. Following digestion, protease and phosphatase inhibitors were added, and the tissue suspension was passed through a 70 μm cell strainer into a 50 mL tube. The filtrate was subjected to sequential centrifugation at 4 °C (300 × g for 10 min, 2,000 × g for 20 min, and 10,000 × g for 30 min). The resulting supernatant was filtered through a 0.22 μm filter and layered onto a 30% sucrose cushion, followed by ultracentrifugation at 100,000 × g for 90 min at 4 °C. The EV pellet was washed with Dulbecco’s phosphate-buffered saline (DPBS) and ultracentrifuged again at 100,000 × g for 90 min at 4 °C. The final pellet was resuspended in 50 μL of ice-cold DPBS for EV characterization and subsequent mass spectrometry analysis.

EV preparations were characterized by negative-stain transmission electron microscopy (TEM) and Western blotting. For TEM, representative fields were imaged at an accelerating voltage of 200 kV and a magnification of 81,000× to assess EV morphology and size. Particle diameters were measured from representative images and summarized for each preparation. For preparations in which multiple particles were measured from an individual animal, the median particle diameter was used for descriptive comparisons between groups. Western blotting was performed to assess the presence of EV-associated proteins, including ALIX, CD9, and TSG101, and the absence of the Golgi-associated protein GM130, which was used as a negative marker to assess potential contamination with non-EV cellular components.

### 2.4 Digestion and mass spectrometry

For each sample, 10 µg protein was buffer-exchanged three times against 25 mM ammonium bicarbonate, dried, resuspended in 50 µL of 25 mM ammonium bicarbonate containing 50% acetonitrile, and digested overnight at 37 °C with 1 µg trypsin and 500 ng Lys-C. Digests were dried and reconstituted in 50 µL of 2% acetonitrile and 0.1% formic acid. Each sample was analyzed by one 5 µL injection followed by a blank cycle.

Peptides were separated on a C18 EasySpray column at 0.350 µL/min and analyzed on a Q Exactive HF-X mass spectrometer. Mobile phase A was 0.1% formic acid in water and B was 0.1% formic acid in acetonitrile. The gradient held 4% B from 0 to 8.75 min, increased to 40% B at 107.5 min, 65% B at 109.5 min and 95% B at 112.5 min, held 95% B until 123.5 min, returned to 4% B at 124.5 min, and ended at 127.5 min. Full MS used resolution 60,000, an AGC target of 3 × 10^6^, maximum injection time 45 ms and m/z 300–1500. Top-20 MS/MS used resolution 30,000, an AGC target of 1 × 10^5^, maximum injection time 54 ms, isolation window 1.3 m/z, normalized collision energy 28 and dynamic exclusion 30 s. Unassigned charges and charge states 1, 7, 8 and >8 were excluded.

### 2.5 Database searching and construction of the analysis matrix

Raw files were processed in Proteome Discoverer 3.3.0 using SEQUEST against the Rattus norvegicus reference proteome UP000002494 with digestion enzymes and contaminant sequences. Dynamic modifications included methionine oxidation, protein N-terminal acetylation, initiator methionine loss, and initiator methionine loss plus N-terminal acetylation. Peptide-spectrum matches were assessed with Percolator. Abundances were normalized using Specific Protein Amount with the rat reference database.

The combined export contained 1,404 protein groups. Its Full Filter flag retained 978 entries after excluding flagged contaminants, keratins, immunoglobulins and entries with fewer than three unique peptides. One retained entry, Q02SZ7, was annotated as bacterial lysyl endopeptidase and was excluded as an exogenous digestion-enzyme entry. The resulting set comprised 977 rat protein groups, including 953 with positive normalized abundances in all nine samples. These 953 groups formed the primary complete-case matrix; the remaining 24 were retained in the filtering audit. No missing values were imputed. Missing gene symbols were filled for display from explicit GN annotations in the source description while retaining accessions as the analysis identifiers.

Quantitative analyses used the nine normalized-abundance columns in the combined LFQ workbook, rather than its vendor-generated ratio columns or the individual identification lists. Columns F1-F3 corresponded to LPS, F4-F6 to Mi, and F7-F9 to SD. Individual sample exports were used only for identification-depth checks. Protein groups, rather than presumed unique genes, were the unit of feature-level analysis.

### 2.6 Pairwise testing and quality assessment

Normalized abundances were log2-transformed. Two-sided Welch t-tests compared LPS with SD, Mi with LPS, and Mi with SD. Benjamini-Hochberg adjustment was performed separately across 953 tests within each contrast [4]. We report nominal P values, adjusted q values, log2 fold changes and unadjusted 95% Welch confidence intervals. Fold changes are ratios of geometric means. Exploratory candidates required nominal P < 0.05 and |log2 fold change| ≥ log2(1.5), equivalent to ≥1.5-fold increase or ≤2/3-fold abundance. This candidate screen does not control the false-discovery rate.

Within-group coefficients of variation (CVs) were calculated from the normalized linear abundances as sample standard deviation divided by mean. Pearson correlations used log2 abundances. Principal-component analysis treated the nine preparations as observations and the 953 protein groups as variables; each protein was centered across samples without variance scaling. Distances between group means were reported as root-mean-square differences across all 953 log2 features, not as distances restricted to the first two principal components. Approximate Hedges g values used the pooled log2 standard deviation and a small-sample correction factor of 0.8; they were treated as descriptive effect-size estimates.

### 2.7 Descriptive treatment trajectories and whole profile distance

For each protein, let a denote mean log2 abundance in LPS minus SD and b denote mean log2 abundance in Mi minus LPS. The residual Mi-versus-SD difference is a + b. Opposite-direction movement was defined by a × b < 0. A stronger descriptive criterion, closer to control, required |a + b| < |a|. For the LPS-screened subset, the signed recovery fraction was defined as follows:

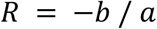

Values 0 < R < 1 indicate movement toward control without crossing it; R = 1 indicates equality of observed group means; 1 < R < 2 indicates overshoot that remains closer to control; and R ≥ 2 indicates overshoot without improved proximity. These labels describe the observed means and do not establish statistical equivalence, efficacy or mechanism. Selection on the LPS contrast and reuse of the LPS mean in both contrasts can favor apparent reversal through regression to the mean.

We assessed the complete profile using a distance-based recovery measure across all 953 proteins. With a and b now denoting the corresponding vectors of protein-level contrasts:

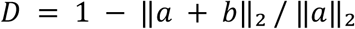

D = 1 indicates coincidence of the treatment and control mean profiles, D = 0 indicates equal distances of LPS and treatment from control, and D < 0 indicates a larger treatment-to-control distance. This measure was not clipped. An exploratory percentile interval was calculated from 10,000 bootstrap resamples of animals within each group (three draws with replacement per group; random seed 20261003). Entire sample columns were resampled to preserve dependence among proteins. The 953-feature set remained fixed. With only three animals per group, this interval describes resampling uncertainty in this cohort and is not a definitive population-level test.

### 2.8 Sensitivity and annotation analyses

Each animal was omitted in turn in nine sensitivity analyses. For each contrast we recorded the range and consistency of fold-change direction; no animal was excluded from the primary analysis. Alternative preprocessing analyses retained proteins with at least two observed abundances in each compared group, or additionally median-centered each sample on the log2 scale. A separate audit reproduced the source filter with Q02SZ7 retained. As a test-family sensitivity, BH adjustment was also applied jointly to all 2,859 pairwise tests. This alternative controls a different family and need not be more conservative for every contrast.

The broad Biological Process, Cellular Component and Molecular Function labels provided in the export were assessed by one-sided hypergeometric overrepresentation tests. The background was all 953 quantified protein groups. We tested LPS-increased candidates, LPS-decreased candidates, and LPS candidates with a nominally significant opposite treatment effect. Labels assigned to fewer than five background proteins and labels beginning with “other” were omitted. BH adjustment covered all 81 annotation tests. These analyses summarize broad supplied annotations and do not establish specific pathway activation or a physically connected protein network.

## 3 Results

### 3.1 Characterization and quantitative data coverage

TEM showed vesicle-like profiles in each group, and the supplied immunoblot panel showed ALIX, CD9 and TSG101 signals (**Figure 1**). Preparation-level median diameters were 87.0, 105.0 and 60.3 nm for SD; 72.0, 68.1 and 72.8 nm for LPS; and 75.9, 63.3 and 70.6 nm for Mi. The corresponding means ± sample SD of the three animal medians were 84.1 ± 22.5, 71.0 ± 2.5 and 69.9 ± 6.3 nm. These descriptive measurements do not establish equivalence of size distributions or exclude differences in particle composition.

**Figure 1.**
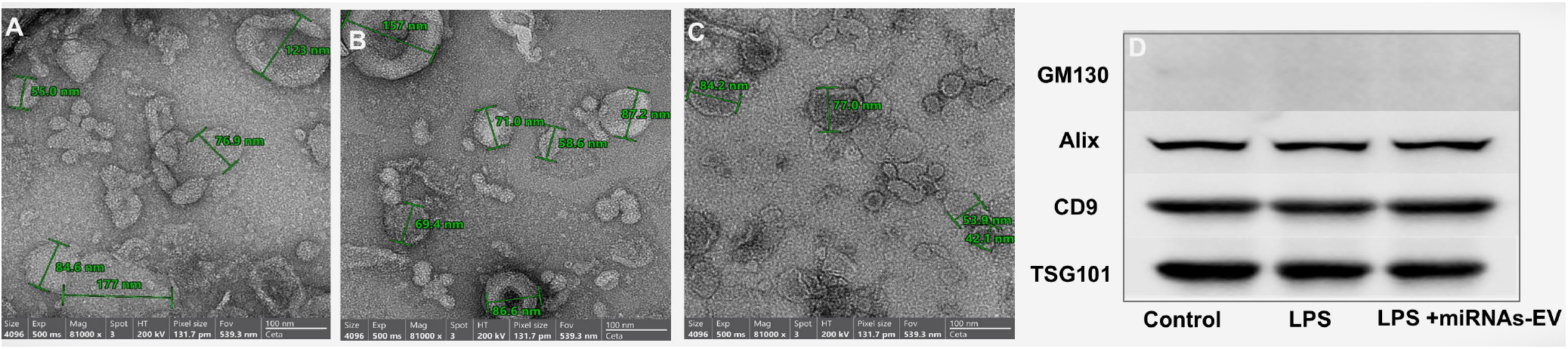
Morphological and immunoblot characterization. (**A-C**) Representative negative-stain TEM fields for brains EVs isolated from SD, LPS and Mi treatment groups respectively. Scale bars and diameter annotations are retained from the supplied source panel. (**D**) Supplied marker-blot panel showing ALIX, CD9 and TSG101 signals and a negative-control target.

After exclusion of Q02SZ7, 977 protein groups remained and 953 were complete across all samples (**Table 1**). Individual identification exports contained 587-986 entries, with the lowest and highest counts both occurring in LPS samples (**Figure 2A**). Individual identification depth and abundance completeness in a combined analysis are different quantities and were not treated as interchangeable. Log2 within-group correlations ranged from 0.948 to 0.990 (**Figure 2B**). Median protein CVs were 18.6% in SD, 25.5% in LPS and 13.3% in Mi. These values summarize combined biological and analytical variation; they do not isolate technical precision. The filtered quantitative matrix contained Cd9, Pdcd6ip/ALIX, Flot1 and Hspa8, together with Canx, Calr, Hspa5 and Hsp90b1. Tsg101 was not found by accession-description or gene-name search in the supplied LFQ export, despite its reported immunoblot signal.

**Table 1.**
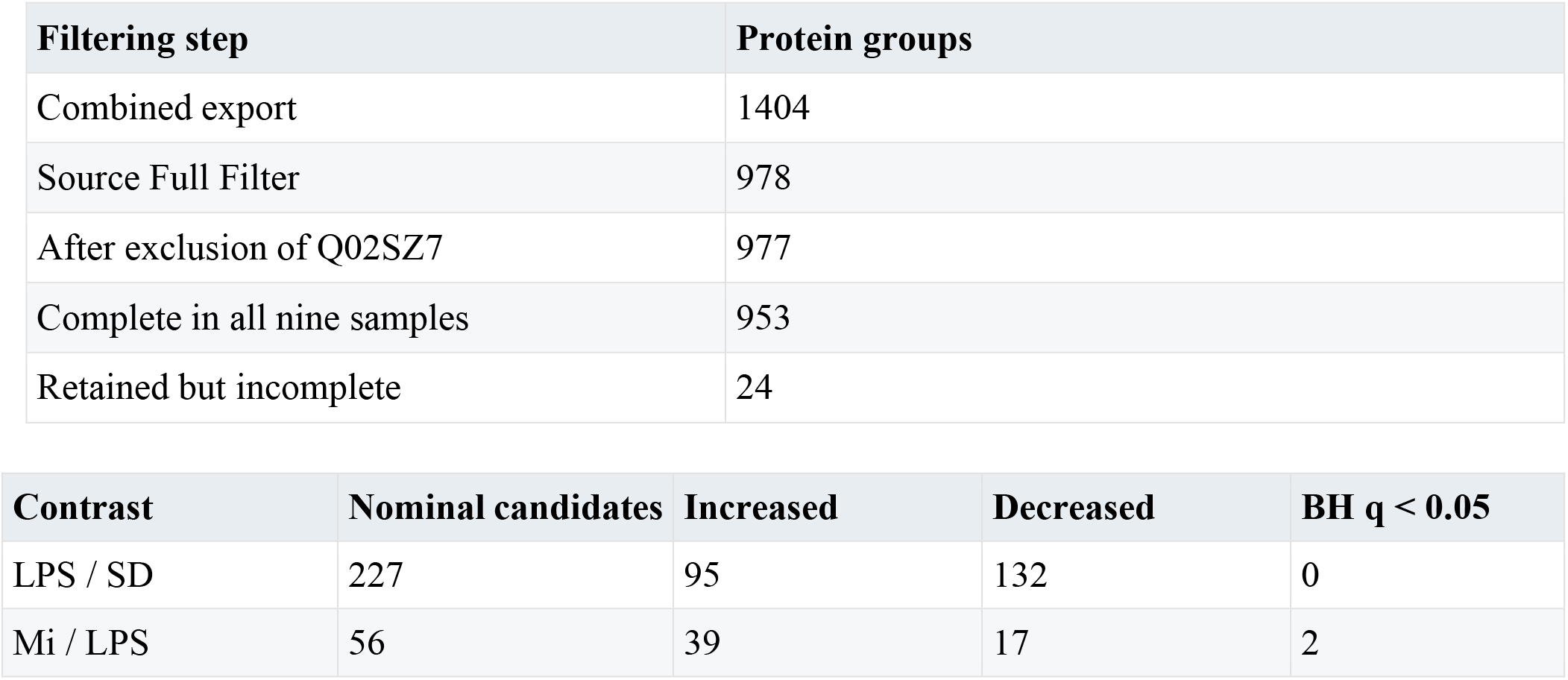
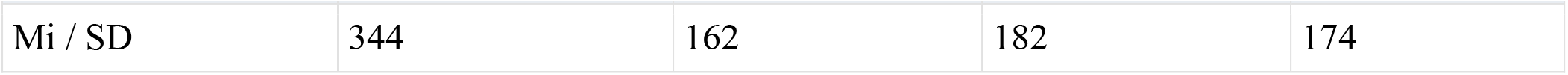
Filtering and pairwise protein-level results. Nominal candidates require P < 0.05 and |log2FC| ≥ log2(1.5). BH adjustment is separate within each 953-protein contrast.

| Filtering step | Protein groups |
| --- | --- |
| Combined export | 1404 |
| Source Full Filter | 978 |
| After exclusion of Q02SZ7 | 977 |
| Complete in all nine samples | 953 |
| Retained but incomplete | 24 |

| Contrast | Nominal candidates | Increased | Decreased | BH q < 0.05 |
| --- | --- | --- | --- | --- |
| LPS / SD | 227 | 95 | 132 | 0 |
| Mi / LPS | 56 | 39 | 17 | 2 |

| Contrast | Nominal candidates | Increased | Decreased | BH $q < 0.05$ |
| --- | --- | --- | --- | --- |
| Mi / SD | 344 | 162 | 182 | 174 |

**Figure 2.**
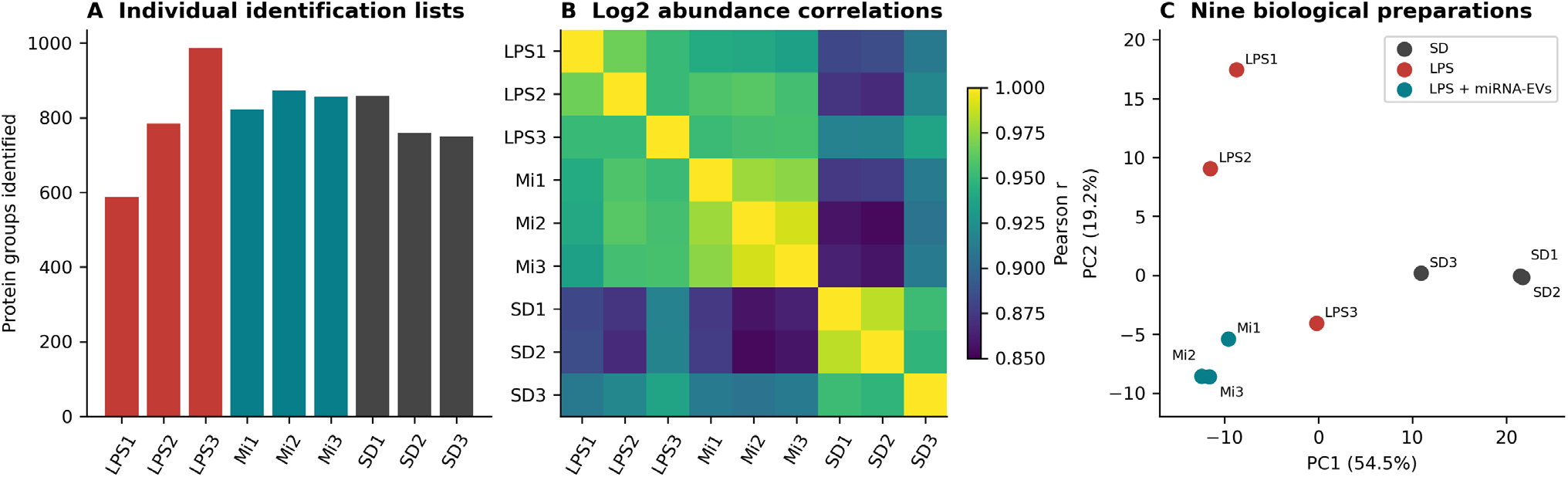
Identification depth and sample-level variation. (**A**) Protein-group counts from the nine individual identification exports, before the combined-analysis filter. (**B**) Pearson correlation matrix of the 953 complete log2-normalized abundance profiles. (**C**) PCA with one point per biological preparation (n = 9), using protein-centered data without variance scaling. PC1 and PC2 explain 54.5% and 19.2% of variance. Mi denotes LPS plus anti-inflammatory miRNA-associated EVs. Numerical replicate labels follow the within-group export order.

Mi denotes LPS plus miRNA-associated EV formulation. Counts refer to protein groups. The 174 Mi/SD FDR findings include 163 that also meet the fold-change cutoff.

### 3.2 LPS associated changes were bidirectional and exploratory

For LPS versus SD, 251 proteins had nominal P < 0.05 and 498 met the 1.5-fold magnitude criterion. Their intersection contained 227 candidates: 95 increased and 132 decreased (**Figure 3A**; **Table 1**). None passed within-contrast BH correction at q < 0.05; the smallest q value was 0.0696. This result limits protein-specific claims and does not justify treating the nominal list as a validated set.

**Figure 3.**
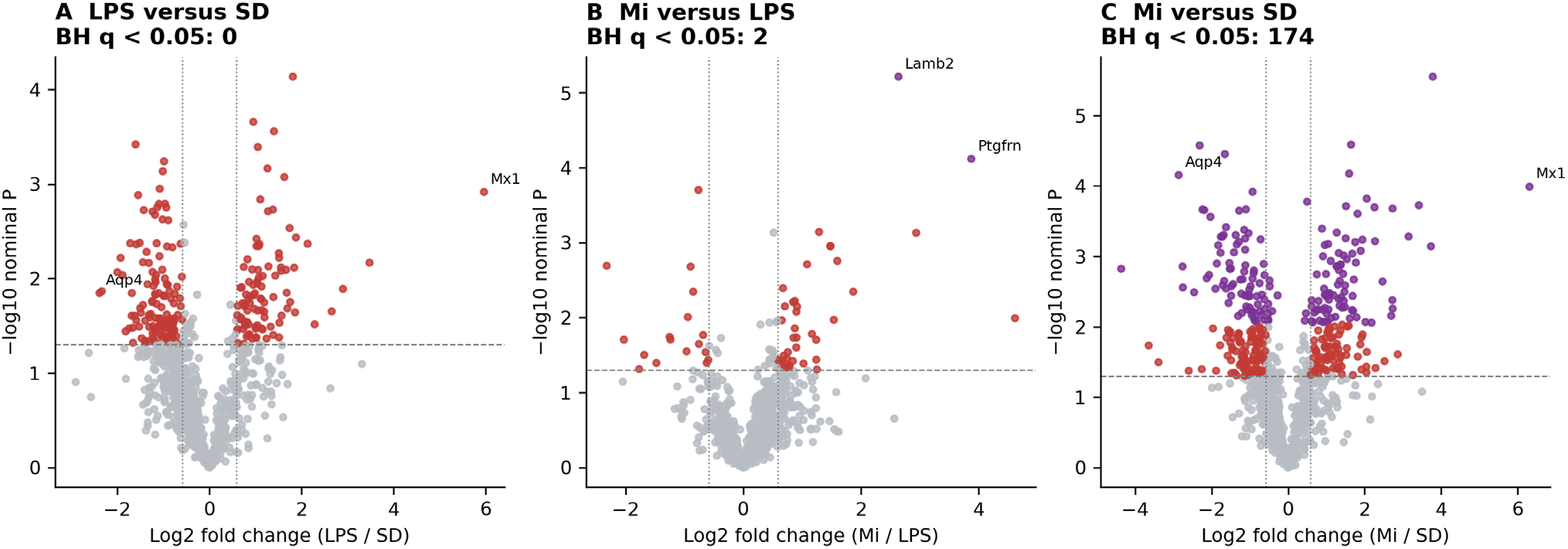
All three differential-abundance contrasts. Each point represents one of 953 protein groups. Purple indicates within-contrast BH q < 0.05. Red indicates the nominal P < 0.05 and 1.5-fold screen without FDR significance; gray indicates other proteins. Horizontal lines mark nominal P = 0.05 and vertical lines mark ±log2(1.5). Fold changes derive from mean log2 abundances. The numbers of FDR findings are 0 for LPS/SD, 2 for Mi/LPS and 174 for Mi/SD.

Mx1 had a 62.5-fold observed increase relative to SD. Aqp4 and Kcnj13 had relative abundances of 0.191 and 0.271, respectively. Kcnj13 encodes Kir7.1 and should not be identified as the astroglial Kir4.1 channel, which is encoded by Kcnj10 [5,6]. Of the 95 increased candidates, 33 had a translational-apparatus annotation, compared with 53 of 953 background proteins (6.25-fold enrichment; annotation q = 1.18 × 10^−20^). Of the 132 decreased candidates, 69 had a plasma-membrane annotation, compared with 310 of 953 background proteins (1.61-fold enrichment; annotation q = 6.53 × 10^−6^). These results support broad compositional patterns within the screened lists.

### 3.3 The treatment profile remained different from control

PC1 and PC2 explained 54.5% and 19.2% of total variance, respectively, in the centered, unscaled PCA (**Figure 2C**). Across the complete feature space, the root-mean-square centroid distance was 0.553 log2 units between Mi and LPS, 0.984 between Mi and SD, and 0.864 between LPS and SD. Thus, the treatment mean profile was closer to LPS than to control and was not globally closer to control than the LPS profile was.

Mi versus LPS yielded 56 nominal candidates under the combined P-value and fold-change screen. Two protein groups passed BH q < 0.05: Lamb2 (6.23-fold; P = 6.14 × 10^−6^; q = 0.00585) and Ptgfrn (14.71-fold; P = 7.65 × 10^−5^; q = 0.0365) (Figure 3B). Both were quantified in all nine samples, so these are increased abundances rather than newly detected proteins. Mi versus SD yielded 174 proteins at q < 0.05, of which 163 also met the 1.5-fold criterion (**Figure 3**). This contrast documents residual differences at the 24 h endpoint.

### 3.4 Selected candidates moved toward control

Among the 227 LPS-screened candidates, 120 (52.9%) changed in the opposite direction after anti-inflammatory miRNAs-EVs +LPS treatment, and 119 (52.4%) were closer to control on the log2 scale. Of the opposite-direction group, 114 approached control without overshoot, five overshot but remained closer, and one overshot without improved proximity. Thirty-nine candidates had R ≥ 0.5. Twenty-three also had nominal P < 0.05 for the opposite treatment effect, and 18 of these had observed treatment-to-control fold changes within the interval from 2/3 to 1.5. Membership in this interval is not an equivalence test.

C3, Wfs1, Tmx1, Agpat3, Cds2, Atp2a2, Atp2a3, Syt1 and Dpysl5 were among these 23 candidates (**Table 2**; **Figure 4**). For example, Wfs1 changed from 0.329-fold of SD under LPS to 0.919-fold after treatment; C3 changed from 2.05-fold to 1.07-fold. All nine retained the opposite-direction pattern in every leave-one-sample-out analysis. However, none had a treatment-versus-LPS q < 0.05. Their consistency support targeted follow-up, while their adjusted results require continued classification as exploratory candidates.

**Table 2.**
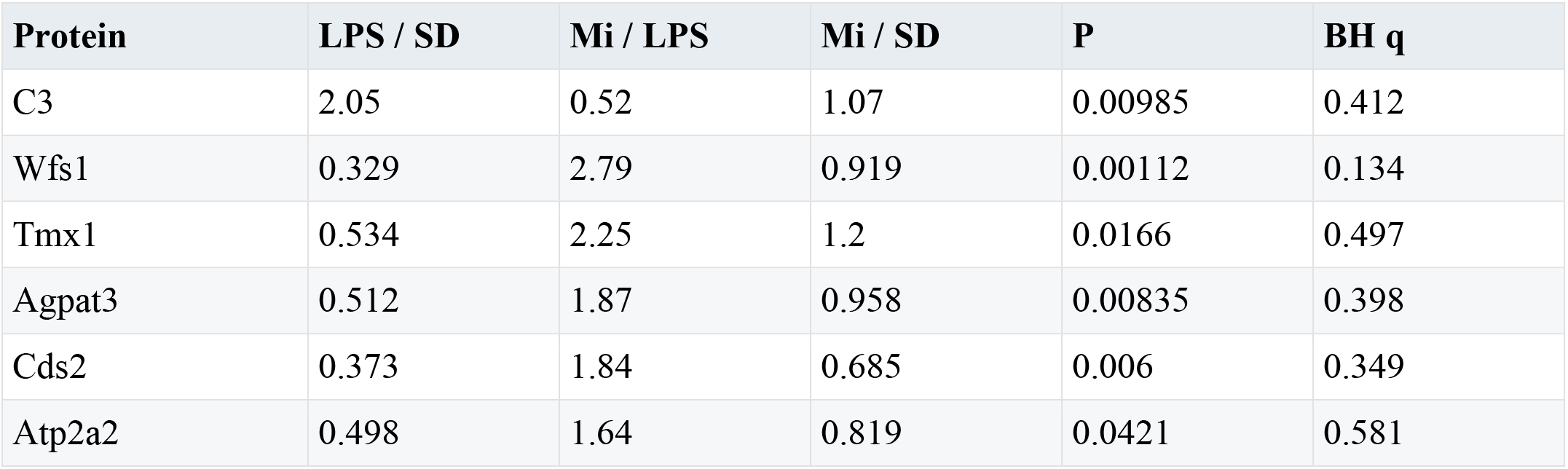
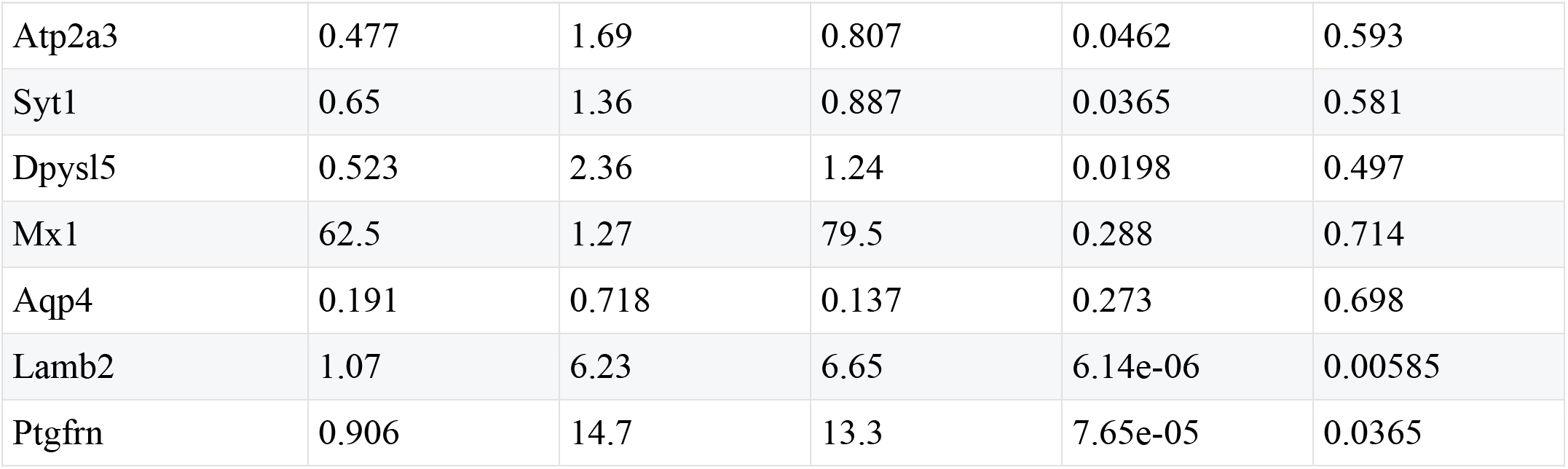
Selected observed protein responses. Fold changes are ratios of geometric means. P and q refer to Mi versus LPS; q is adjusted across 953 proteins. All values are exploratory estimates from three preparations per group.

| Protein | LPS / SD | Mi / LPS | Mi / SD | P | BH q |
| --- | --- | --- | --- | --- | --- |
| C3 | 2.05 | 0.52 | 1.07 | 0.00985 | 0.412 |
| Wfs1 | 0.329 | 2.79 | 0.919 | 0.00112 | 0.134 |
| Tmx1 | 0.534 | 2.25 | 1.2 | 0.0166 | 0.497 |
| Agpat3 | 0.512 | 1.87 | 0.958 | 0.00835 | 0.398 |
| Cds2 | 0.373 | 1.84 | 0.685 | 0.006 | 0.349 |
| Atp2a2 | 0.498 | 1.64 | 0.819 | 0.0421 | 0.581 |
| Atp2a3 | 0.477 | 1.69 | 0.807 | 0.0462 | 0.593 |
| Syt1 | 0.65 | 1.36 | 0.887 | 0.0365 | 0.581 |
| Dpysl5 | 0.523 | 2.36 | 1.24 | 0.0198 | 0.497 |
| Mx1 | 62.5 | 1.27 | 79.5 | 0.288 | 0.714 |
| Aqp4 | 0.191 | 0.718 | 0.137 | 0.273 | 0.698 |
| Lamb2 | 1.07 | 6.23 | 6.65 | 6.14e-06 | 0.00585 |
| Ptgfrn | 0.906 | 14.7 | 13.3 | 7.65e-05 | 0.0365 |

**Figure 4.**
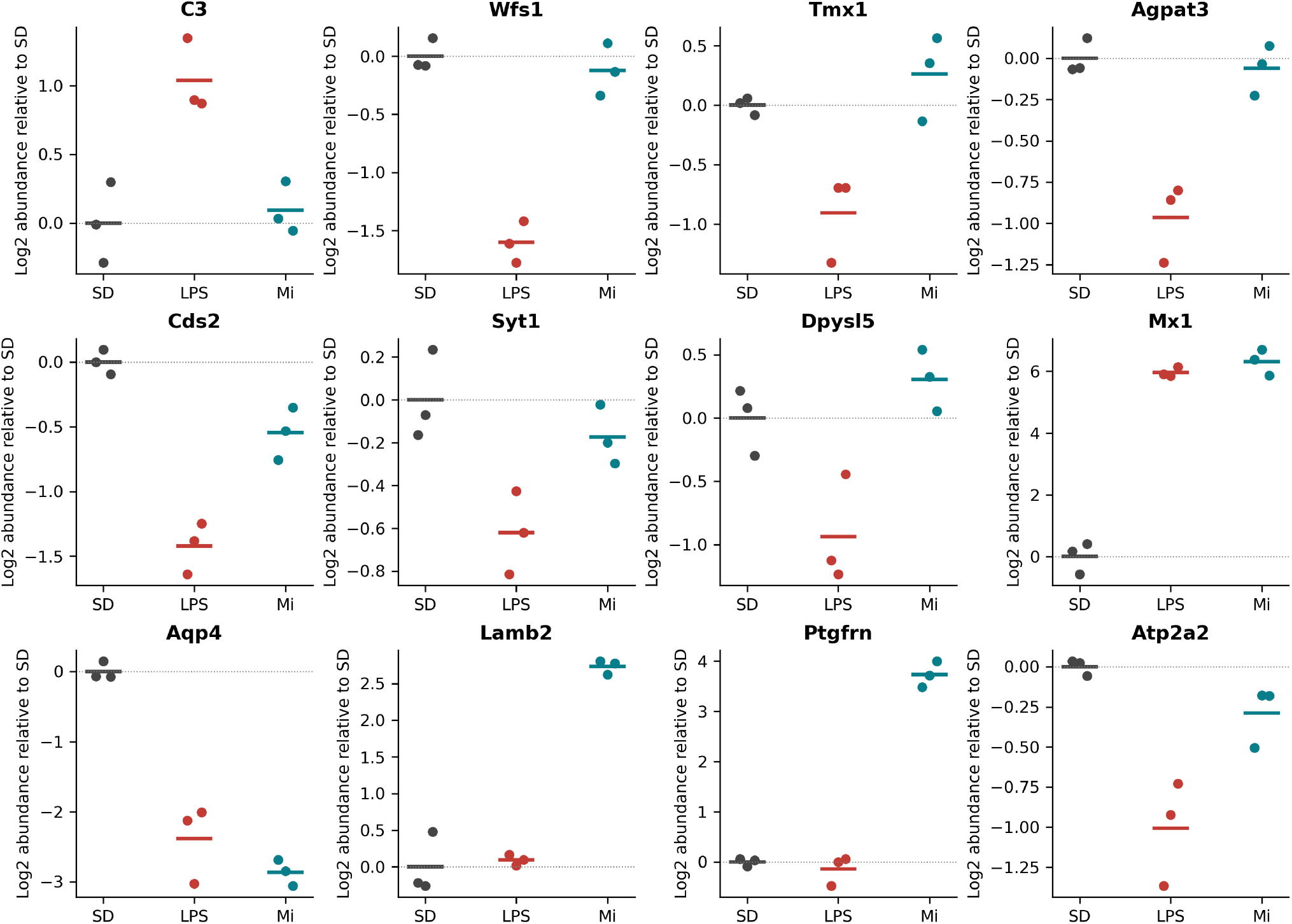
Individual observations for selected response candidates. Each point is one independent preparation; horizontal segments indicate group means. Values are log2 abundances centered to the SD mean for that protein. SD, vehicle control; LPS, LPS only; Mi, LPS plus formulation. Panels have separate vertical scales. The figure displays observed trajectories rather than validated rescue. Atp2a3 and the complete candidate list are included in Table 2 and the workbook.

Mx1 remained high after treatment (79.5-fold relative to SD), while Aqp4 remained low (0.137-fold). The direct Mi-versus-LPS differences for these proteins were not significant even at nominal P < 0.05. The observations therefore support persistent deviations from control, rather than demonstrated treatment-induced worsening of either protein.

A treatment mean near the SD mean does not establish equivalence. For the nine proposed recovery candidates, treatment q values range from 0.134 to 0.593.

### 3.5 Sensitivity analyses limited claims of global recovery

The complete-profile distance measure was D = -0.139, with an exploratory animal-bootstrap 95% interval of -0.307 to 0.069 (**Figure 5**). Its leave-one-sample out range was -0.198 to 0.028. The point estimate indicates greater treatment-to-control distance, but the uncertainty does not distinguish this reliably from equal distance or modest convergence. It should not be interpreted as evidence that the treatment worsened tissue function.

**Figure 5.**
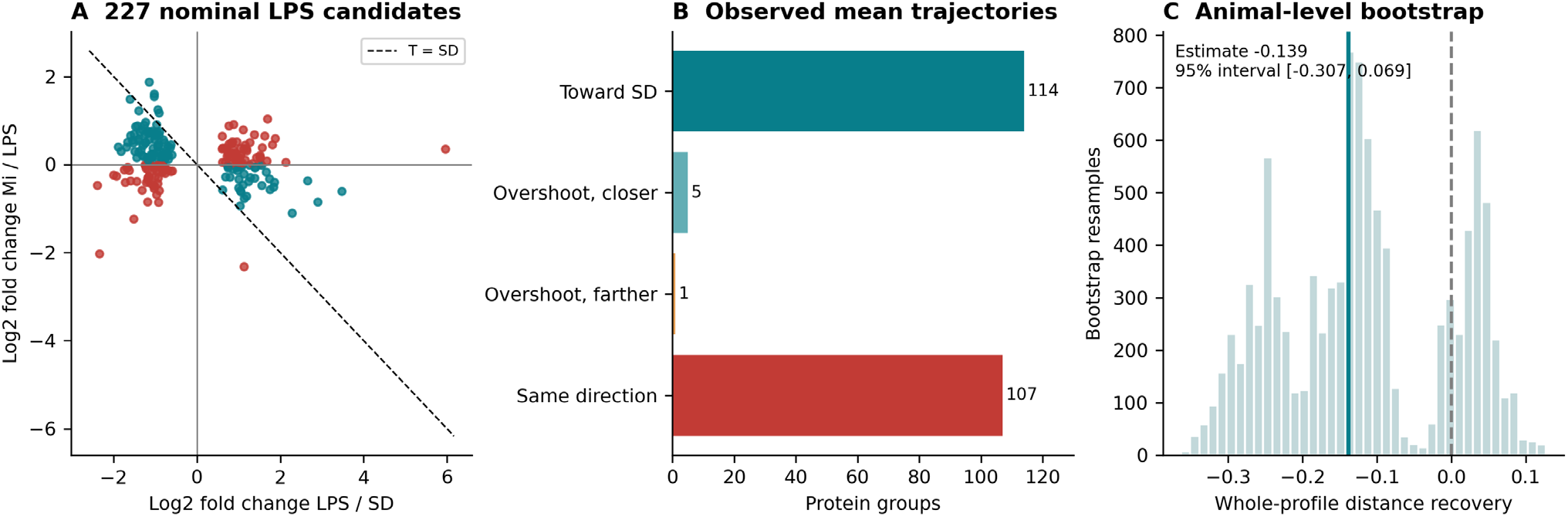
Descriptive trajectories and whole-profile distance. (A) LPS/SD and Mi/LPS log2 contrasts for 227 nominal LPS-screened candidates. Teal indicates a treatment mean closer to SD; red indicates no improved proximity. The dashed diagonal indicates equal observed treatment and SD means. (B) Deterministic trajectory counts: 114 toward control without overshoot, five overshooting but closer, one overshooting without improved proximity, and 107 continuing in the same direction or unchanged. (C) Distribution of the distance-recovery measure from 10,000 animal-level bootstrap resamples within groups. The observed value is −0.139 and the percentile 95% interval is −0.307 to 0.069. Negative values indicate a larger treatment-to-control distance. This small-cohort resampling interval is exploratory.

Of the 120 opposite-direction LPS candidates, 92 retained that direction in all nine omission analyses. Allowing at least two observed samples per compared group produced 233 nominal LPS-versus-SD candidates, with no FDR-significant proteins in that contrast, and preserved two treatment-versus-LPS FDR findings. Additional sample-median centering changed the corresponding counts to 223 nominal LPS candidates and one FDR finding in each of the LPS-versus-SD and treatment-versus-LPS contrasts. Under median centering, Ptgfrn retained treatment-versus-LPS significance (q = 0.0342), whereas Lamb2 did not (q = 0.165). These changes show that some threshold decisions depend on preprocessing. The primary analysis therefore retained the laboratory-supplied normalization.

## 4 Discussion

The main finding is selective remodeling of the protein composition of brain EV-enriched preparations after exposure to the miRNA-associated formulation. At 24 h, several proteins had observed abundances closer to control, while the overall treatment profile remained distinct from control. Lamb2 and Ptgfrn provided the strongest direct treatment-versus-LPS evidence under the primary analysis. The other highlighted responses were nominal candidates whose directions were stable to single-sample omission, but whose treatment contrasts did not survive proteome-wide FDR correction. Together, these observations motivate a model in which inflammatory signaling, organelle homeostasis and membrane trafficking influence the recovered extracellular protein profile. Figure 6 connects this model to the measured findings while separating established biology from causal steps that remain untested in this experiment.

**Figure 6.**
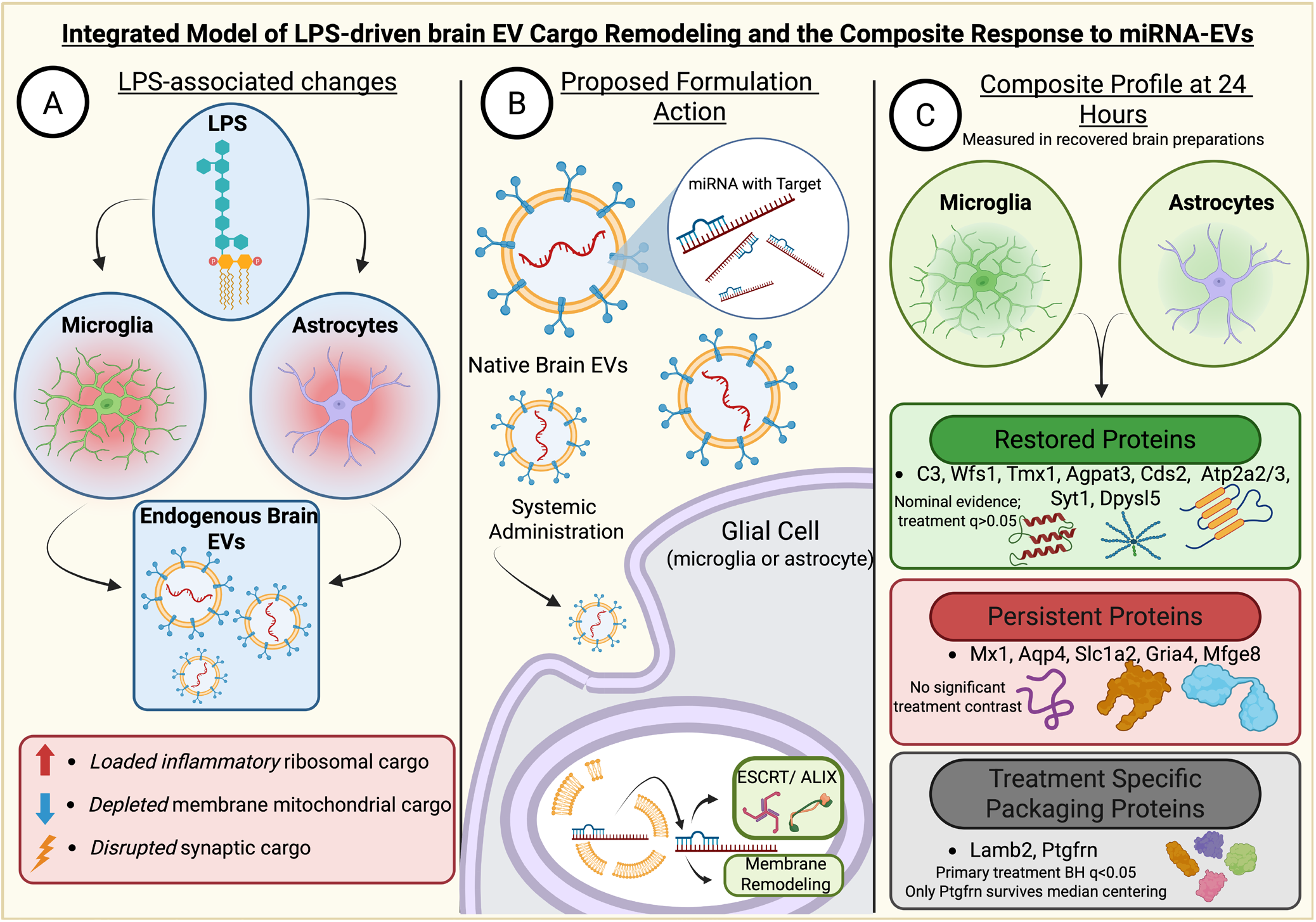
Integrated model of LPS-associated brain EV protein changes and responses to miRNA-associated EVs. (A) LPS-associated changes include increased inflammatory/ribosomal candidates, decreased membrane-associated candidates and altered synaptic-associated proteins. Microglia and astrocytes are plausible sources based on prior studies; cellular origin and selective packaging were not established [1,2,7,9]. (B) MiRNA-associated EVs prepared from naïve rat brain are administered systemically. Glial uptake, RNA delivery and regulation of cargo selection are proposed mechanisms; ESCRT/ALIX and membrane remodeling are candidate processes, not demonstrated miRNA targets [20,21]. MiRNA localization and target engagement remain unverified, and administered material may contribute to the recovered brain EV-enriched preparation. (C) Three response patterns coexist at 24 h: toward-control changes in C3, Wfs1, Tmx1, Agpat3, Cds2, Atp2a2/3, Syt1 and Dpysl5 (treatment-versus-LPS BH q > 0.05); residual deviations in Mx1, Aqp4, Slc1a2, Gria4 and Mfge8, without significant treatment-versus-LPS differences; and FDR-significant treatment-associated increases in Lamb2 and Ptgfrn. Slc1a2 was not identified in the initial nominal LPS screen, and only Ptgfrn remained significant after additional median centering. These patterns do not establish functional recovery, distinct EV subpopulations or treatment-specific sorting. The overall proteome showed no demonstrated convergence toward controls. Dashed arrows and question marks indicate hypotheses; cartoons do not establish colocalization. n = 3 preparations per group. SD, vehicle control; ICV, intracerebroventricular; EV, extracellular vesicle; ESCRT, endosomal sorting complex required for transport; BH, Benjamini–Hochberg; FDR, false discovery rate.

An LPS challenge can engage both inflammatory and interferon-related signaling. Genetic disruption experiments established that TLR4 responses involve MyD88 and TRIF, with TRIF contributing to IRF3 activation and interferon-β production [7]. This provides a plausible upstream context for the large observed increase in Mx1, an interferon-induced GTP-binding protein [8]. Mx1 remained elevated in both LPS and Mi preparations, at 62.5-fold and 79.5-fold of control, respectively. Because the direct treatment comparison was not significant, these data support a residual interferon-associated protein signature rather than treatment-induced enhancement of this pathway. Neither TLR4 activation nor the downstream transcriptional response was measured directly; extracellular Mx1 abundance cannot establish the activity or cellular location of either process.

The broad annotation patterns further suggest that LPS changes the kinds of material recovered in the EV-enriched fraction. Translational-apparatus annotations were enriched among increased candidates, whereas plasma-membrane annotations were enriched among decreased candidates. These patterns could arise from altered source-cell states, vesicle subtype proportions, protein sorting, release, clearance or tissue-derived material that co-isolates with EVs. They do not demonstrate increased translation within vesicles. State-dependent cargo profiles and inflammatory responses in recipient cells have been demonstrated for experimentally isolated microglial EVs [1], and acute neuroinflammation has been associated with altered cortical EV cargo [2]. Those studies make an EV-related contribution plausible, but they do not identify the source cells or the active particles in the present mixed brain preparation.

C3 offers a biologically relevant link between the inflammatory environment and possible effects on neural tissue. Its observed abundance changed from 2.05-fold of control under LPS to 1.07-fold after formulation exposure. In other experimental systems, activated microglia induce a C3-positive astrocyte response through combinations of inflammatory mediators [9], and complement perturbation modifies microglial synapse elimination in Alzheimer mouse models [10]. These findings support testing complement-related consequences here, but they cannot establish that the C3 measured in our preparations originated from astrocytes or mediated synaptic injury. Total C3 protein also does not distinguish intact precursor from complement activation products. The nominal treatment response therefore identifies a candidate change in complement-associated material, not demonstrated inhibition of complement activity.

The persistence of high Mx1 and low Aqp4 provides a useful counterpoint to the C3 trajectory. Different inflammatory and membrane-associated responses may have distinct time courses or different sensitivities to the formulation. Alternatively, these proteins may report different source-cell or particle populations. Aqp4 abundance in an isolated fraction cannot establish water-channel function, perivascular localization, edema resolution or astrocyte number in the tissue. Moreover, C3, Mx1 and Aqp4 need not reside in the same vesicles. The separation of their responses argues for measuring multiple biological processes independently rather than assigning the preparation a single inflammatory or recovery state. Cell-resolved localization, complement-fragment assays and functional measurements would distinguish these possibilities.

The additional residual-deviation candidates Slc1a2, Gria4 and Mfge8 strengthen the description of a mixed treatment-associated state. Their Mi/SD abundances were 0.465-, 0.535- and 0.317-fold, with BH q values of 0.0119, 0.0237 and 0.00840, respectively. None had a significant direct Mi-versus-LPS contrast, even at nominal P < 0.05. Slc1a2 also did not meet the initial nominal LPS screen (P = 0.0652), so its inclusion in Figure 6 describes a residual deviation relative to control rather than a statistically established LPS lesion that treatment failed to reverse. These proteins should therefore be treated as a descriptive response group, not as a validated persistent pathway or a measure of treatment resistance.

Wfs1, Tmx1, Atp2a2 and Atp2a3 nominate ER homeostasis and calcium handling as a focused mechanistic hypothesis. Wfs1 rose from 0.329-fold of control in LPS preparations to 0.919-fold after treatment. WFS1 can restrain ATF6α signaling by promoting HRD1-dependent ubiquitination and degradation of ATF6α in cellular models [11]. The present trajectory is therefore compatible with a change in material associated with ER stress regulation. It does not establish that WFS1 activity was restored inside brain cells, that ATF6 signaling decreased, or that the relevant WFS1 pool was delivered by EVs. These alternatives require matched tissue or cell measurements of WFS1, ATF6 processing and downstream stress responses.

The concurrent trajectories of Tmx1 and the calcium pumps make this hypothesis more specific without making it causal. TMX1 interacts with SERCA2b at ER–mitochondria contact sites and can inhibit SERCA activity under the conditions examined in cultured cancer cells, thereby changing calcium partitioning and mitochondrial metabolism [12]. Thus, an increase in TMX1 cannot simply be equated with improved calcium pumping. ATP2A2 protein-group abundance also does not resolve the SERCA2b isoform without isoform-specific peptide evidence. The observed approach toward control of Atp2a2 and Atp2a3, together with the modest overshoot of Tmx1, could reflect altered ER membrane representation, compensatory regulation or changed protein export. The informative next step is to measure ER calcium content, stimulus-evoked mitochondrial calcium uptake and respiration in the same biological system. Their relationships to the EV profile would test whether the proteomic pattern accompanies functional organelle recovery.

Agpat3 and Cds2 provide a biochemical connection between organelle composition and membrane trafficking. AGPAT3, also termed LPAAT3, contributes to phosphatidic-acid production; experimental manipulation of this enzyme changes Golgi membrane tubulation, organization and trafficking [13]. CDS2 converts phosphatidic acid into CDP-diacylglycerol for phosphatidylinositol synthesis. Its deletion in macrophages revealed compensatory lipid metabolism and impaired maintenance of the phosphatidylinositol cycle during sustained stimulation [14]. These studies caution against interpreting enzyme abundance as a direct measure of lipid flux. Here, Agpat3 approached the control mean, whereas Cds2 showed only partial convergence. This pattern motivates targeted lipidomics of matched tissue and EV-enriched fractions, including phosphatidic acid and phosphatidylinositol species, rather than a claim that membrane lipid composition or EV biogenesis was restored.

The Syt1 and Dpysl5 trajectories extend the candidate set to neural membrane and cytoskeletal-associated material. SYT1 has an experimentally established role in fast calcium-triggered transmitter release at central synapses [15], but a change in extracellular SYT1 abundance is not a measurement of neurotransmission. Greater recovery of neural proteins could reflect selective packaging, a shift in the fraction of neuron-derived particles, or release of membrane material during tissue injury and processing. These alternatives can produce similar proteomic patterns with very different biological meanings. Colocalization with validated particle markers, comparison with matched tissue abundance and direct assays of synaptic activity would be needed before assigning a synaptic-protective mechanism.

Ptgfrn and Lamb2 deserve separate emphasis because their treatment-versus-LPS increases met the primary FDR threshold. PTGFRN has been experimentally used as an EV-enriched scaffold for surface presentation of engineered cargo [16]. Its 14.71-fold increase here makes it a candidate marker of formulation-associated membrane remodeling or altered particle representation. It does not demonstrate activation of the same sorting mechanism, particularly because the published scaffold work involved engineered producer cells and cargo fusions. PTGFRN was present in all nine preparations, and its contribution could derive from administered carrier material, endogenous particles, or changes in recovery of PTGFRN-containing membranes. Its persistence after additional median normalization supports prioritization for independent validation.

Lamb2 increased 6.23-fold under the primary treatment comparison, but did not retain FDR significance after additional median centering. Laminin β2 can organize presynaptic calcium-channel clusters and active zones at the neuromuscular junction [17]. That observation establishes a specific extracellular matrix–synapse interaction in another anatomical setting; it does not show that increased LAMB2 in a brain EV-enriched preparation repairs central synapses. Extracellular matrix association, vascular or other source-cell contributions, and carrier-associated material remain plausible explanations. Together, the Ptgfrn and Lamb2 findings illustrate why a reproducible treatment response need not represent reversal of an LPS-induced defect. Their biological meaning depends on localization, particle association and function, not only on the size of the fold change.

**Figure 6** proposes a conditional mechanism: formulation components reach responsive cells, biologically available miRNA engages intracellular silencing machinery, and altered gene regulation changes inflammatory or organelle states, with secondary effects on the released or retained EV-associated proteome. This sequence would connect a molecular intervention to the mixed extracellular protein response. However, each intermediate remains to be established. Uptake of labeled material alone would not demonstrate cytosolic RNA delivery, Argonaute loading or repression of a relevant target. The proteomic experiment did not measure sequence-specific target engagement, so no particular miRNA–target interaction or direct regulation of Wfs1, C3, Ptgfrn or another candidate can yet be assigned. Proteins that increase could reflect indirect responses even if the initiating RNA acts through target repression.

The formulation method introduces a further mechanistic uncertainty because miRNA may be intraluminal, surface-associated or associated with the loading reagent. Quantitative studies of native EV preparations have found low average miRNA copy numbers [18], and experiments in defined cell systems have shown that EV binding need not lead to effective miRNA delivery or target regulation [19]. These findings do not determine the loading or activity of this engineered preparation; they show why those properties must be measured for the actual formulation. Carrier components, residual free RNA or reagent-associated RNA could also contribute. A second explanation requires no reprogramming of endogenous secretion: persistence or retention of administered material, together with altered clearance or co-isolation, could change the recovered protein profile. The figure therefore presents these as competing routes to the measured endpoint rather than treating EV sorting as an established mechanism.

Endosomal cargo selection is one candidate connection between these cellular responses and the extracellular profile. The syndecan-syntenin-ALIX system can support intraluminal budding, with syntenin binding ALIX and engaging ESCRT-associated machinery [20]. Perturbation studies also show that individual ESCRT components and ALIX can affect EV composition and release differently across cellular systems [21]. This makes ESCRT/ALIX-dependent sorting a plausible checkpoint to test, but does not show that the administered miRNAs regulate it. In the present matrix, Pdcd6ip/ALIX itself had a Mi/LPS ratio of 0.937 (P = 0.511; q = 0.835). Its unchanged total abundance neither demonstrates nor excludes a change in recruitment, localization or activity. Establishing the proposed route would require sequence-specific target engagement together with imaging or biochemical measurements of endosomal recruitment and a perturbation experiment linking that process to the extracellular protein response.

The aggregate results constrain how the mechanistic model should be interpreted. Although 120 of 227 nominal LPS candidates moved in the opposite direction, only 119 were actually closer to control. The nine highlighted candidates retained their direction in all omission analyses, but none passed the primary treatment FDR threshold. In contrast, 174 proteins differed between Mi and control at q < 0.05, and the whole-profile distance measure was −0.139 with an exploratory 95% interval of −0.307 to 0.069. These findings describe selective convergence embedded within a broader treatment-associated state. They neither establish global normalization nor exclude a functional benefit that was not measured. A biologically useful response need not reproduce every control abundance, but that possibility must be supported by functional outcomes rather than inferred from the proteomic distance.

Two statistical features are especially relevant to this interpretation. First, the LPS mean enters the LPS-versus-control and treatment-versus-LPS contrasts with opposite signs, allowing sampling error to generate apparent reversal; selecting the initial LPS-responsive list adds regression-to-the-mean bias. Second, three animals per group remain three biological replicates regardless of the number of quantified proteins. Correlated protein responses cannot substitute for independent animals, and omission analyses assess sensitivity within this cohort rather than external reproducibility. Equal-protein loading and laboratory normalization also make these compositional measurements: they do not quantify secretion per brain or copies of cargo per vesicle. These considerations favor effect estimates, individual observations and prespecified validation over probabilistic claims of pathway restoration.

## 5. Limitations and Future Directions

This study examined a small male-only cohort at a single endpoint, and the extent to which the observed responses vary with sex and time remains to be established. Although the proteomic findings identify candidate biological pathways, the causal relationships proposed in **Figure 6** require direct mechanistic validation.

Future studies should extend these findings to independent cohorts incorporating both sexes and multiple time points. Targeted proteomics and complementary measurements in matched brain tissue should prioritize treatment-associated proteins Ptgfrn and Lamb2, candidates showing shifts toward control abundance such as C3, Wfs1 and Agpat3, and persistent deviations involving Mx1 and Aqp4. Mechanistic investigations should test whether sequence-specific miRNA target engagement influences endosomal cargo selection and membrane remodeling. Integrating these analyses with measurements of complement activity, ER stress, calcium handling, lipid composition and synaptic function would clarify the biological consequences of selective EV cargo changes and determine whether they precede sustained functional improvement.

## 6. Conclusion

Reanalysis of 953 complete protein groups identified 227 nominal LPS-associated candidates and two FDR-significant treatment-versus-LPS abundance increases, Lamb2 and Ptgfrn. Selected candidates moved toward control, while treatment-to-control comparisons and whole-profile distances did not demonstrate global normalization. The data supports an exploratory map of formulation-associated protein changes in brain EV-enriched preparations and a focused panel for independent validation. They do not yet establish anti-inflammatory miRNA-specific therapeutic rescue or reprogramming of endogenous EV sorting. The candidate responses nominate a testable connection between inflammatory state, ER calcium handling and membrane composition, with cellular target engagement and particle provenance as key unresolved steps.

## References

1. Santiago JV, Natu A, Ramelow CC, Rayaprolu S, Xiao H, Kumar V, Kumar P, Seyfried NT, Rangaraju S. Identification of State-Specific Proteomic and Transcriptomic Signatures of Microglia-Derived Extracellular Vesicles. Mol Cell Proteomics. 2023 Dec;22(12):100678. doi: 10.1016/j.mcpro.2023.100678. Epub 2023 Nov 11. PMID: 37952696; PMCID: PMC10755493.

2. Vassileff N, Spiers JG, Juliani J, Lowe RGT, Datta KK, Hill AF. Acute neuroinflammation promotes a metabolic shift that alters extracellular vesicle cargo in the mouse brain cortex. J Extracell Biol. 2024 Jun 27;3(7):e165. doi: 10.1002/jex2.165. Erratum in: J Extracell Biol. 2025 Jul 24;4(7):e70070. doi: 10.1002/jex2.70070. PMID: 38947878; PMCID: PMC11212288.

3. Welsh JA, Goberdhan DCI, O’Driscoll L, Buzas EI, Blenkiron C, Bussolati B, Cai H, Di Vizio D, Driedonks TAP, Erdbrügger U, Falcon-Perez JM, Fu QL, Hill AF, Lenassi M, Lim SK, Mahoney MG, Mohanty S, Möller A, Nieuwland R, Ochiya T, Sahoo S, Torrecilhas AC, Zheng L, Zijlstra A, Abuelreich S, Bagabas R, Bergese P, Bridges EM, Brucale M, Burger D, Carney RP, Cocucci E, Crescitelli R, Hanser E, Harris AL, Haughey NJ, Hendrix A, Ivanov AR, Jovanovic-Talisman T, Kruh-Garcia NA, Ku’ulei-Lyn Faustino V, Kyburz D, Lässer C, Lennon KM, Lötvall J, Maddox AL, Martens-Uzunova ES, Mizenko RR, Newman LA, Ridolfi A, Rohde E, Rojalin T, Rowland A, Saftics A, Sandau US, Saugstad JA, Shekari F, Swift S, Ter-Ovanesyan D, Tosar JP, Useckaite Z, Valle F, Varga Z, van der Pol E, van Herwijnen MJC, Wauben MHM, Wehman AM, Williams S, Zendrini A, Zimmerman AJ; MISEV Consortium; Théry C, Witwer KW. Minimal information for studies of extracellular vesicles (MISEV2023): From basic to advanced approaches. J Extracell Vesicles. 2024 Feb;13(2):e12404. doi: 10.1002/jev2.12404. Erratum in: J Extracell Vesicles. 2024 May;13(5):e12451. doi: 10.1002/jev2.12451. PMID: 38326288; PMCID: PMC10850029.

4. Yoav Benjamini, Yosef Hochberg, Controlling the False Discovery Rate: A Practical and Powerful Approach to Multiple Testing, Journal of the Royal Statistical Society: Series B (Methodological), Volume 57, Issue 1, January 1995, Pages 289–300.

5. National Center for Biotechnology Information. Kcnj13 potassium inwardly-rectifying channel, subfamily J, member 13, Rattus norvegicus. Gene ID 94341. https://www.ncbi.nlm.nih.gov/gene/94341. Accessed October 3, 2026.

6. National Center for Biotechnology Information. Kcnj10 potassium inwardly-rectifying channel, subfamily J, member 10, Rattus norvegicus. Gene ID 29718. https://www.ncbi.nlm.nih.gov/gene/29718. Accessed October 3, 2026.

7. Yamamoto M, Sato S, Hemmi H, Hoshino K, Kaisho T, Sanjo H, Takeuchi O, Sugiyama M, Okabe M, Takeda K, Akira S. Role of adaptor TRIF in the MyD88-independent toll-like receptor signaling pathway. Science. 2003 Aug 1;301(5633):640–3. doi: 10.1126/science.1087262. Epub 2003 Jul 10. PMID: 12855817.

8. National Center for Biotechnology Information. Mx1 MX dynamin like GTPase 1, Rattus norvegicus. Gene ID 24575. https://www.ncbi.nlm.nih.gov/gene/24575. Accessed October 3, 2026.

9. Liddelow SA, Guttenplan KA, Clarke LE, Bennett FC, Bohlen CJ, Schirmer L, Bennett ML, Münch AE, Chung WS, Peterson TC, Wilton DK, Frouin A, Napier BA, Panicker N, Kumar M, Buckwalter MS, Rowitch DH, Dawson VL, Dawson TM, Stevens B, Barres BA. Neurotoxic reactive astrocytes are induced by activated microglia. Nature. 2017 Jan 26;541(7638):481–487. doi: 10.1038/nature21029. Epub 2017 Jan 18. PMID: 28099414; PMCID: PMC5404890.

10. Hong S, Beja-Glasser VF, Nfonoyim BM, Frouin A, Li S, Ramakrishnan S, Merry KM, Shi Q, Rosenthal A, Barres BA, Lemere CA, Selkoe DJ, Stevens B. Complement and microglia mediate early synapse loss in Alzheimer mouse models. Science. 2016 May 6;352(6286):712–716. doi: 10.1126/science.aad8373. Epub 2016 Mar 31. PMID: 27033548; PMCID: PMC5094372.

11. Fonseca SG, Ishigaki S, Oslowski CM, Lu S, Lipson KL, Ghosh R, Hayashi E, Ishihara H, Oka Y, Permutt MA, Urano F. Wolfram syndrome 1 gene negatively regulates ER stress signaling in rodent and human cells. J Clin Invest. 2010 Mar;120(3):744–55. doi: 10.1172/JCI39678. Epub 2010 Feb 15. PMID: 20160352; PMCID: PMC2827948.

12. Raturi A, Gutiérrez T, Ortiz-Sandoval C, Ruangkittisakul A, Herrera-Cruz MS, Rockley JP, Gesson K, Ourdev D, Lou PH, Lucchinetti E, Tahbaz N, Zaugg M, Baksh S, Ballanyi K, Simmen T. TMX1 determines cancer cell metabolism as a thiol-based modulator of ER-mitochondria Ca2+ flux. J Cell Biol. 2016 Aug 15;214(4):433–44. doi: 10.1083/jcb.201512077. Epub 2016 Aug 8. PMID: 27502484; PMCID: PMC4987292.

13. Schmidt JA, Brown WJ. Lysophosphatidic acid acyltransferase 3 regulates Golgi complex structure and function. J Cell Biol. 2009 Jul 27;186(2):211–8. doi: 10.1083/jcb.200904147. PMID: 19635840; PMCID: PMC2717635.

14. Collins DM, Janardan V, Barneda D, Anderson KE, Niewczas I, Taylor D, Qiu D, Jessen HJ, Lopez-Clavijo AF, Walker S, Raghu P, Clark J, Stephens LR, Hawkins PT. CDS2 expression regulates de novo phosphatidic acid synthesis. Biochem J. 2024 Oct 16;481(20):1449–1473. doi: 10.1042/BCJ20240456. PMID: 39312194; PMCID: PMC11555650.

15. Geppert M, Goda Y, Hammer RE, Li C, Rosahl TW, Stevens CF, Südhof TC. Synaptotagmin I: a major Ca2+ sensor for transmitter release at a central synapse. Cell. 1994 Nov 18;79(4):717–27. doi: 10.1016/0092-8674(94)90556-8. PMID: 7954835.

16. Dooley K, McConnell RE, Xu K, Lewis ND, Haupt S, Youniss MR, Martin S, Sia CL, McCoy C, Moniz RJ, Burenkova O, Sanchez-Salazar J, Jang SC, Choi B, Harrison RA, Houde D, Burzyn D, Leng C, Kirwin K, Ross NL, Finn JD, Gaidukov L, Economides KD, Estes S, Thornton JE, Kulman JD, Sathyanarayanan S, Williams DE. A versatile platform for generating engineered extracellular vesicles with defined therapeutic properties. Mol Ther. 2021 May 5;29(5):1729–1743. doi: 10.1016/j.ymthe.2021.01.020. Epub 2021 Jan 21. PMID: 33484965; PMCID: PMC8116569.

17. Nishimune H, Sanes JR, Carlson SS. A synaptic laminin-calcium channel interaction organizes active zones in motor nerve terminals. Nature. 2004 Dec 2;432(7017):580–7. doi: 10.1038/nature03112. PMID: 15577901.

18. Chevillet JR, Kang Q, Ruf IK, Briggs HA, Vojtech LN, Hughes SM, Cheng HH, Arroyo JD, Meredith EK, Gallichotte EN, Pogosova-Agadjanyan EL, Morrissey C, Stirewalt DL, Hladik F, Yu EY, Higano CS, Tewari M. Quantitative and stoichiometric analysis of the microRNA content of exosomes. Proc Natl Acad Sci U S A. 2014 Oct 14;111(41):14888–93. doi: 10.1073/pnas.1408301111. Epub 2014 Sep 29. PMID: 25267620; PMCID: PMC4205618.

19. Albanese M, Chen YA, Hüls C, Gärtner K, Tagawa T, Mejias-Perez E, Keppler OT, Göbel C, Zeidler R, Shein M, Schütz AK, Hammerschmidt W. MicroRNAs are minor constituents of extracellular vesicles that are rarely delivered to target cells. PLoS Genet. 2021 Dec 6;17(12):e1009951. doi: 10.1371/journal.pgen.1009951. PMID: 34871319; PMCID: PMC8675925.

20. Baietti MF, Zhang Z, Mortier E, Melchior A, Degeest G, Geeraerts A, Ivarsson Y, Depoortere F, Coomans C, Vermeiren E, Zimmermann P, David G. Syndecan-syntenin-ALIX regulates the biogenesis of exosomes. Nat Cell Biol. 2012 Jun 3;14(7):677–85. doi: 10.1038/ncb2502. PMID: 22660413.

21. Colombo M, Moita C, van Niel G, Kowal J, Vigneron J, Benaroch P, Manel N, Moita LF, Théry C, Raposo G. Analysis of ESCRT functions in exosome biogenesis, composition and secretion highlights the heterogeneity of extracellular vesicles. J Cell Sci. 2013 Dec 15;126(Pt 24):5553–65. doi: 10.1242/jcs.128868. Epub 2013 Oct 8. PMID: 24105262.

22. Yan X, Chen X, Shan Z, Bi L. Engineering Exosomes to Specifically Target the Mitochondria of Brain Cells. ACS Omega. 2023 Dec 11;8(51):48984–48993. doi: 10.1021/acsomega.3c06617. PMID: 38162779; PMCID: PMC10753542.

